# Morphological hallmarks of dopaminergic neurodegeneration are associated with altered neuron function in *Caenorhabditis elegans*

**DOI:** 10.1101/2023.08.22.554364

**Authors:** Andrew S. Clark, Javier Huayta, Katherine S. Morton, Joel N. Meyer, Adriana San-Miguel

## Abstract

*Caenorhabditis elegans* (*C. elegans*) is an excellent model system to study neurodegenerative diseases, such as Parkinson’s disease, as it enables analysis of both neuron morphology and function in live animals. Multiple structural changes in neurons, such as cephalic dendrite morphological abnormalities, have been considered hallmarks of neurodegeneration in this model, but their relevance to changes in neuron function are not entirely clear. We sought to test whether hallmark morphological changes associated with chemically induced dopaminergic neuron degeneration, such as dendrite blebbing, breakage, and loss, are indicative of neuronal malfunction and result in changes in behavior. We adapted an established dopaminergic neuronal function assay by measuring paralysis in the presence of exogenous dopamine, which revealed clear differences between *cat-2* dopamine deficient mutants, wildtype worms, and *dat-1* dopamine abundant mutants. Next, we integrated an automated image processing algorithm and a microfluidic device to segregate worm populations by their cephalic dendrite morphologies. We show that nematodes with dopaminergic dendrite degeneration markers, such as blebbing or breakage, paralyze at higher rates in a dopamine solution, providing evidence that dopaminergic neurodegeneration morphologies are correlated with functional neuronal outputs.

## 1. Introduction

Neurodegenerative diseases, such as Parkinson’s disease (PD) increasingly plague the aging population (Dorsey et al., 2018). The cause(s) of PD is still unknown, yet it is thought to result from a combination of genetic and environmental factors (Calahorro & Ruiz-Rubio, 2011). The hallmark of the disease is the progressive loss of dopamine (DA) neurons in the *substantia nigra* in the brain, which ultimately presents itself as uncontrollable movement (Calahorro & Ruiz-Rubio, 2011; Wooten, 1997). Due to the mainly idiopathic nature of PD, it has been hypothesized that environmental exposures may play a key contributing role in the onset of the disease. In fact, there is strong evidence that specific environmental pesticides, such as rotenone and paraquat, are linked to increased PD susceptibility (Gasser, 2015; Nandipati & Litvan, 2016).

*C. elegans* is widely used as a PD model due to its simplicity and versatility for culturing, imaging, and functional behavior studies. These nematodes have 302 neurons, eight of which are dopaminergic (Brenner, 1974; Youssef et al., 2019). Several DA-mediated behaviors have been well documented, such as basal slowing, odor avoidance, and swimming-induced paralysis (Allen et al., 2011; Boyd et al., 2012; Hart, 2006; McDonald et al., 2007; Smith et al., 2019). Analysis of dopamine signaling in *C. elegans* has been aided by loss of function mutations in genes such as *cat-2* (a tyrosine hydroxylase), (Sawin et al., 2000), which leads to DA deficiency; and *dat-1* (a dopamine reuptake transporter), which leads to extracellular DA abundance(Hardaway et al., 2015; Kudumala et al., 2019). DA-deficient worms fail to slow their locomotion when placed on food, which is a common phenotype for healthy, wildtype worms (Sawin et al., 2000). This behavioral assay, known as basal slowing, is commonly used to test for DA dysfunction (Smith et al., 2019). In contrast, due to the inability to clear excess DA, *dat-1* worms quickly exhibit a paralysis phenotype when swimming in water, which is known as swimming-induced paralysis (SWIP) (Hardaway et al., 2015; McDonald et al., 2007). This phenotype is associated with increased dopamine in the synaptic cleft, and thus animals with normal or reduced levels of dopamine do not exhibit obvious SWIP phenotypes (Smith et al., 2019).

Various chemical exposures can induce dopaminergic neuron degeneration (Alam & Schmidt, 2002; Coulom & Birman, 2004; Mello et al., 2022; Norazit et al., 2010; Offenburger et al., 2018; Offenburger & Gartner, 2018; Tanner et al., 2011; Testa et al., 2005; Uversky, 2004). 6-hydroxydopamine (6-OHDA) is a chemical commonly used to measure the effects of DA dysfunction due to its specificity for DA neurons (Glinka et al., 1997; Omura et al., 2012; Smith et al., 2019). The morphological effects of 6-OHDA exposures have been well characterized and have shown to induce clearly detectable neurodegeneration in the cephalic (CEP) neurons (Bijwadia et al., 2021; Cothren et al., 2018; González-Hunt et al., 2014; Nass et al., 2002; Smith et al., 2019). The CEP dendrites in healthy nematodes are nearly straight lines with no apparent irregularities, particularly early in life. However, the CEP dendrites in worms exposed to 6-OHDA often exhibit clear breakage (the disappearance of dendrite) or blebbing (protrusions from the string-like dendrites) (Bijwadia et al., 2021; Clark et al., 2023; González-Hunt et al., 2014). To gain a better understanding of chemical-induced morphological changes in DA neurons, levels of dendritic degeneration have been categorized into a variety of classifications, from presence/absence to more nuanced scales such as 5- or 7-point systems (Bijwadia et al., 2021; Cothren et al., 2018).

However, there have been very limited efforts to test whether morphological changes correlate with behavioral changes in exposed animals. Smith et al. connected DA morphology to behavior by comparing the neuron morphology and worm exploration areas associated with various levels of 6-OHDA exposure (Smith et al., 2019); however, in that study, population-level morphological and behavioral metrics were compared, in different populations.

In this study, we take advantage of unbiased quantitative DA degeneration image analysis and microfluidics-based sorting to segregate *individual* animals by their dendrite morphologies and assess if these are associated with changes in dopaminergic function. We first aimed to develop a behavioral assay capable of distinguishing healthy, DA-deficient, and DA-abundant worms. We accomplished this by adapting the SWIP assay (Hardaway et al., 2015; McDonald et al., 2007) to include small levels of exogenous DA (DA-SWIP). Traditionally, wild type worms do not paralyze at high rates during the SWIP assay, and modifications that are detected in this phenotype are the result of endogenous dopamine accumulation, resulting in above-background SWIP rates. By adding exogenous dopamine, the dopamine transport and signaling system can be evaluated in worms that have decreased dopamine levels. DA-SWIP thus examines alterations in a phenotype clearly present in controls rather than the emergence of a phenotype. Addition of exogenous DA resulted in paralysis of both wildtype and *cat-2* worms, with *cat-2* worms paralyzing at a slower rate. Using this assay, we reveal that worms exposed to 50 mM 6-OHDA paralyze quicker in DA than control worms. Moreover, we show that worms exposed to 50 mM 6-OHDA also exhibit an altered basal slowing response, supporting that DA-SWIP can be used to study dopaminergic function. We then tested whether worms with specific dendritic morphologies exhibited differences in functional behavior. In our previous study, we developed an automated image processing algorithm capable of rapidly detecting and quantifying levels of dopaminergic neurodegeneration from maximum intensity projections (MIP) (Clark et al., 2023). Thus, we integrated our previously developed algorithm, fluorescent microscopy, and a microfluidic device to create a semi-automated sorting platform. Using this platform, we split nematode populations exposed to the same chemical and environmental conditions into three groups based on their dopaminergic dendrite morphologies. Our results show that worms with healthy dendrites paralyze less than worms with dendritic blebs and breakage, thus showing that morphological neurodegeneration markers, such as blebs and breaks, are linked with neuronal dysfunction.

## 2. Materials and methods

### a. Worm strains and culture

Worms were cultured on nematode growth medium (NGM) seeded with *E*. coli OP50 lawns at 20 °C. The dopamine swimming induced paralysis experiments used BY200 [vtIs1 (*dat*-1p::GFP, *rol*-6)] as the wildtype strain, RM2702 [*dat-1*(ok157)] as the dopamine abundant strain, and CB1112 [*cat-2*(e1112)] as the dopamine deficient strain. The locomotion rate experiments also used CB1141 [*cat-4* (e1141) V]. Sorting experiments exclusively used the BY200 [vtIs1 (*dat*-1p::GFP, *rol*-6)] strain. were provided by the *Caenorhabditis* Genetics Center, which is funded by NIH Office of Research Infrastructure Programs (P40 OD010440).

### b. 6-OHDA exposure

Prior to behavioral analysis, all worm populations were age synchronized by extracting eggs from gravid adult worms using standard 1% bleach and 0.1 M NaOH bleaching solutions (Stiernagle, 2006). Eggs were washed three times using M9 buffer with 0.01% Triton TX100. Eggs were left in the M9 buffer for 16 hours for all worms to reach L1 arrest. L1 worms were then placed on seeded NGM plates until they became L4 larvae, about 48 hours. At 48 hours, worms were washed 3 times with M9 and placed in a 600 µL solution of either 50 mM 6-OHDA or 0 mM vehicle control for 1 hour. 50 mM 6-OHDA exposure were made from a stock solution of 100 mM 6-OHDA (Sigma-Aldrich) in 20 mM L-Ascorbic Acid (AA). The 0 mM vehicle control worms were placed in 20 mM AA. After the 1-hour exposure, worms were washed 5 times to completely remove all 6-OHDA. In-between washes, worms were allowed to settle to the bottom of the microcentrifuge tube through gravity. After washing, worms were plated on new, seeded NGM plates and permitted to recover overnight.

### c. Dopamine swimming induced paralysis

A solution of 300 µM dopamine hydrochloride (Sigma-Aldrich) in deionized water was made from a stock solution of 30 mM dopamine. 50 µL of the dopamine solution was pipetted into one well of a 96-well plate. Worms were then picked from a plate with food into the dopamine solution. After removal of the worms from the pick, a timer was started, and the 96-well plate was placed under the dissecting microscope. Worms were immediately measured for paralysis. Thus, t = 0 starts after the time it takes to move the well plate from the side of the bench to under the microscope (about 10-15 seconds). Worms were marked as moving if they were able to complete one full undulatory movement. Movement rates were measured either every 5 minutes up to 25 minutes or every minute up to 10 minutes. Due to the limited number of available worms in the sorting experiments, 10-minute videos were taken for each technical replicate. Since videos required more prep work, t = 0 is 30 seconds after worms were placed in the dopamine solution. DA-SWIP video files were blinded prior to analysis for paralysis rates.

### d. Locomotion rate

Assay plates were prepared as previously described (Sawin et al., 2000). Briefly, K-agar plates were prepared by spreading concentrated HB101 *E. coli* bacteria (OD >6.0) to form a ring with approximately 1 cm inner diameter and 3 cm outer diameter. Plates were left overnight at 20 °C. Plates with no bacteria were also left overnight at 20 °C. BY200 animals were age-synchronized by bleaching gravid adults and placing recovered eggs in K+ media overnight (Stiernagle, 2006). L1 larval stage animals were transferred to K-agar plates containing OP50 *E. coli* bacteria and grown for 48 hours to reach the L4 larval stage. L4 animals were exposed to 0, 10, 25, or 50 mM 6-OHDA (Sigma-Aldrich) and 10mM ascorbic acid for 1 hour (Hartman et al., 2019). Animals were then washed three times with K media and transferred to K-agar plates containing OP50 *E. coli* bacteria for 24 hours. CB1112 and CB1141 animals were grown in the same manner but no exposure to 6-OHDA was performed. After 24 hours, worms were washed three times with K media and 5-15 worms were transferred in a drop of buffer using a pipette tip to the center of an assay plate. The liquid used for transfer was absorbed using a Kimwipe. After 5 minutes, the animals were recorded in 20 second intervals using a Keyence BZX-2710 microscope. Body bends were counted for each animal, with a total of three biological replicates per treatment.

### e. Microfluidic sorting device and operation

We used a microfluidic sorting device previously developed in our lab (D. Midkiff & San-Miguel, 2019; San-Miguel et al., 2016). This device consists of worm and flush inlets, an imaging channel with an overhead step to improve worm orientation, and two outlets (keep and trash). Worm movement and reagent flow was controlled by a custom pressure box and computer-aided graphical user interface. The microfluidic device was fabricated using standard photo and soft lithography techniques (Duffy et al., 1998; Refai & Blakely, 2019). Briefly, we utilized a 20:1 PDMS elastomer to crosslinker ratio to improve material flexibility and permit on-chip valve operation. A 10:1 layer was used on top as support. The PDMS channels were bonded to a cleaned glass slide using oxygen plasma.

Microfluidic device operation was performed similarly to that described previously without using a cooling chamber for immobilization (San-Miguel et al., 2016). Instead, three plates of young adult worms exposed to 50 mM 6-OHDA at the L4 larval stage were washed three times with 0.01% Triton-X in M9 buffer and then paralyzed with 5 mM tetramisole. The flush inlet consisted of 0.01% Triton-X in M9 with 2 mM tetramisole. Importantly, worms would begin moving in the imaging chamber if the flush inlet did not contain tetramisole. Since air bubbles cause fluid control issues in microfluidic devices, prior to loading worms into the device, all valves and channels were degassed. Valves were filled with a 50% glycerol mixture to match the refractive index of PDMS and decrease glare when imaging.

Worms were loaded into the device using pressure driven flow. When a worm entered the imaging chamber all valves were then closed. If more than one worm entered the imaging chamber, all worms were immediately expelled to the trash outlet. If the worm entered the imaging chamber tail first, the worm was then expelled to the trash outlet. Worms coming in tail first accounted for ∼50% of worms entering the device. If a single worm entered the imaging chamber head first, a z-stack of the CEP neurons would be taken and flattened into a maximum intensity projection (MIP). Subsequently, an adapted version of our previously reported image processing algorithm, AUDDIT (Clark et al., 2023), would analyze and report the median break percent of each dendrite and the total number of blebs. Worms were then either sent to the collection chamber or trash chamber depending on the group being collected at the time. Once ∼30 worms were collected per group, the outlet tubing was cleared by running ∼4 mL of the flush solution through the device. After collecting the required number of worms for each group, worms were then placed on new, seeded NGM plates and allowed to recover overnight (∼16 hours) prior to being analyzed for their functional readout. Three biological replicates were conducted for the sorting experiments on separate days. The sorting and image processing platform was controlled with a custom written application developed with the App Designer on MATLAB 2020b.

### f. Microscopy and imaging

Worms were imaged in the microfluidic device mentioned above. Due to the overhead step in the imaging chamber of the device (**Fig. S1**), ∼90% of worms were in the correct dorso-ventral orientation that allows observation of all four CEP neurons in the MIP image. When a worm entered the imaging chamber headfirst, a z-stack image was taken using 40 stacks with a 1 µm step size and a 60ms exposure. The MIP was the only file saved for downstream analysis of the collected groups. Images were acquired using an inverted Leica DMi8 microscope with a 63x objective (NA = 1.40), a Hamamatsu Orca-Fusion camera, and a metal halide light source.

### g. Statistical analysis

All dopamine paralysis experiments shown were conducted using three biological and three or more technical replicates. Statistical tests were completed using the Statistics and Machine Learning Toolbox on MATLAB 2022b. Paralysis rate comparisons and neurodegeneration metric comparisons were tested with one way ANOVA and a Bonferroni multiple corrections procedure. Differences were deemed significant if the p value was under 0.05. Paralysis rates were also compared using the MatSurv function (Creed et al., 2020), which outputs a Kaplan-Meier curve with a Log-Rank test (**Figs. S2-4**). Replicates from experimental groups were pooled prior to analysis. Statistical analysis for the locomotion rate experiments was performed using two-way ANOVA (MATLAB 2022b) followed by two-tailed Student’s T-test (MS Excel for MS 365). Three biological replicates were used. Sample size numbers for each experiment are included in **Table S1**.

## 3. Results and discussion

### a. Dopamine Swimming Induced Paralysis (DA-SWIP) as a behavioral readout for dopaminergic neuron function

With the goal to develop a readout that can robustly and efficiently measure functional outputs of dopaminergic neurons, we first aimed to differentiate behaviors between wildtypes and DA-deficient or abundant mutants. Since SWIP reveals excess dopamine, we reasoned that such a test would not reveal differences in degenerated neurons that exhibit reduced function. Instead, we developed a new behavioral assay by adapting the well-known SWIP protocol to include 300 µM of dopamine (DA-SWIP). We hypothesized that adding exogenous DA would show differences between the three groups such that DA-abundant *dat-1* worms would paralyze quickly, DA-deficient *cat-2* worms would rarely paralyze, and wildtype worms would exhibit intermediate paralysis rates.

In this assay, worms were picked from a plate with a platinum wire into a small volume of the DA solution and monitored for movement at specific time points under a dissecting microscope. Movement was defined as the worm completing a full sinusoidal body movement within the 10 second observation window. Worms were monitored for movement every 5 minutes up to 25 minutes. After ∼20 minutes, wildtype and *dat-1* strains were nearly fully paralyzed, while approximately 60% of *cat-2* worms remained moving (**Fig. 1A**). Moreover, at earlier time points, a clear difference between all three strains became evident. For example, at 5 minutes, approximately 70%, 50%, and 20% of DA-deficient *cat-2*, wildtype, DA-abundant *dat-1* worms remained moving. Notably, these experiments show the importance of measuring paralysis at multiple time points, as difference between *dat-1* and wildtype worms would not have been revealed if movement was only measured at 20 or 25 minutes.

**Figure 1.**
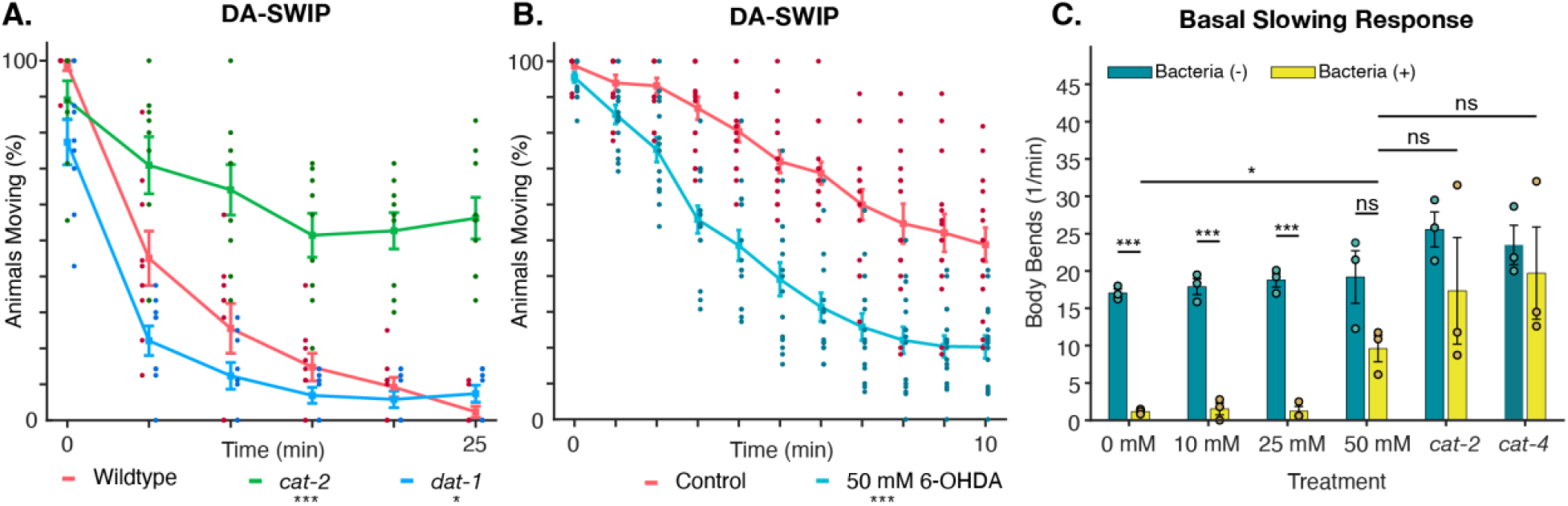
Differences in dopaminergic function, such as dopamine swimming induced paralysis (DA-SWIP) rates and basal slowing responses, are evident in dopaminergic dysfunctional worm strains and worms exposed to 50 mM 6-OHDA. **A**. DA-SWIP paralysis rates of DA-deficient *cat-2* worms (CB1112, green lines), DA-abundant *dat-1* worms (RM2702, blue lines), and wildtype worms (BY200, red lines). **B**. DA paralysis rates of worms exposed to 50 mM 6-OHDA (blue lines) compared to movement rates of worms exposed to the vehicle control (red lines). Significance in curves were determined using the Log-rank test (* p<0.05; *** p<0.001). **C**. Body bends for BY200 (wildtype) animals exposed to 6-OHDA, and positive control strains, CB1112 (*cat-2*) and CB1141 (*cat-4*). Two-way ANOVA with Dunnett’s test, followed with Student’s T-test. p > 0.05 (ns), p ≤ 0.05 (*), p<0.001 (***). Error bars depicted are SEM.

### b. Worms exposed to 6-OHDA exhibit increased paralysis rates in dopamine

Since DA-SWIP revealed significant differences in worm strains with irregular DA levels, we next aimed to test whether worms with 6-OHDA-induced morphological degeneration would exhibit a similar phenotype to either DA-dysfunctional strain. We exposed L4 stage worms to 50 mM 6-OHDA, a dosage previously shown to induce morphological alterations in dopaminergic neurons (Smith et al., 2019), and compared them to a group exposed to vehicle for one hour. After an overnight recovery on plate, we analyzed each group’s dopaminergic function by measuring DA-SWIP rates every minute for 10 minutes (**Fig. 1B**). After ∼2 minutes, it became evident that 6-OHDA worms paralyzed quicker than control worms. To validate that the response was in fact a dopaminergic phenotype, we then compared the basal slowing response of worms exposed to various concentrations of 6-OHDA, unexposed worms, as well as *cat-2* and *cat-4* positive control worm strains (**Fig. 1C**). Worms exposed to 50 mM 6-OHDA showed no statistically significant difference in locomotion between groups with presence or absence of bacteria, indicating a reduction in their basal slowing response. This effect is comparable to the reduction in basal slowing response observed in positive control, defective strains CB1112 (*cat-2*) and CB1141 (*cat-4*), which provides evidence that 6-OHDA exposure induces neuronal dysfunction that can also be revealed by the DA-SWIP response in **Fig. 1B**.

### c. Semi-automated platform to sort worms by level of dopaminergic neuron degeneration

To test whether dendritic morphology is associated with functional changes, we developed a pipeline to sort and collect worms based on their CEP dendrite morphologies. Our approach to accomplish this goal integrated two technologies we have previously: a microfluidic device capable of imaging and sorting worms (D. F. Midkiff et al., 2022; San-Miguel et al., 2016) and an automated image processing algorithm used to detect CEP dendrite morphologies (**Fig. 2A**) (Clark et al., 2023). This semi-automated platform enabled worm imaging, automated dopaminergic dendrite analysis, and worm sorting at a throughput of up to 75 worms per hour.

**Figure 2.**
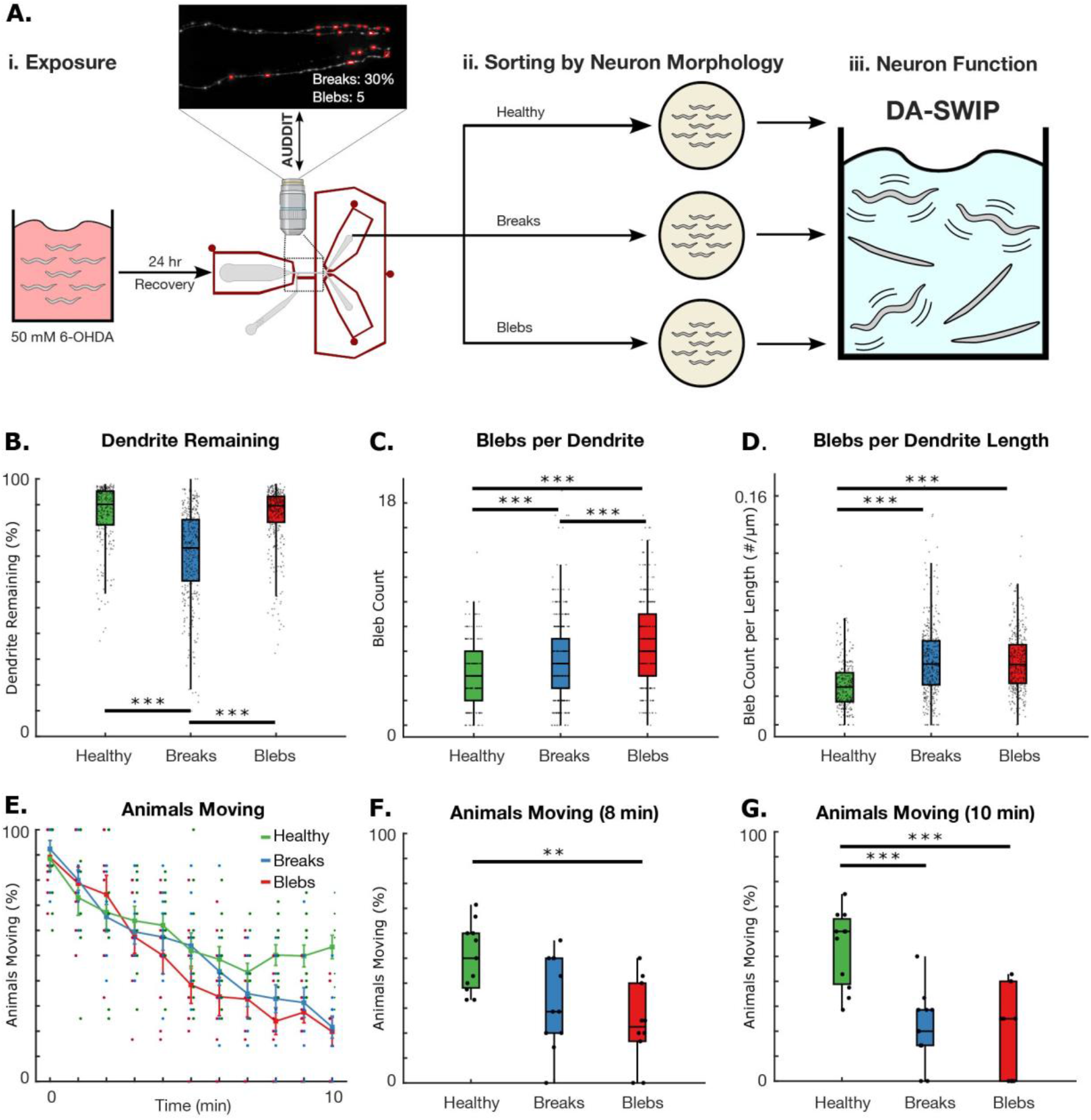
Worms with hallmark morphological neurodegeneration markers have altered dopamine induced swimming paralysis rates. **A**. Experimental pipeline to sort worms exposed to the same chemical exposure by CEP dendrite morphology and test neuron function. **B.-D**. Degeneration metrics obtained from the image processing algorithm for each collected sorted group. **B**. Percentage of dendrite remaining. **C**. Bleb count per dendrite. **D**. Feature count per dendrite length, which considers the total length of the individual dendrite. One-way ANOVA was performed on each group using the Bonferroni multiple correction method. (* p<0.05; ** p<0.01; *** p<0.001). **E**. Percentage of animals moving in 300 µM dopamine with healthy dendrites (green), dendrites with breaks (blue), and dendrites with blebs (red). Log-rank test (p = 0.077). Error bars are SEM. **F-G**. Paralysis rates for each sorted group at the **F**. 8-minute and **G**. 10-minute time points. Statistical significance was determined using one-way ANOVA and the Bonferroni multiple corrections method. (* p<0.05; ** p<0.01; *** p<0.001).

Prior to sorting, we exposed L4 worms to 50 mM 6-OHDA for one hour to obtain a population of worms with varying levels of morphological neurodegeneration. After an overnight recovery, worms were immobilized and loaded into the microfluidic sorting device. After worms individually entered the imaging chamber, epifluorescence z-stack images were acquired of fluorescently labeled CEP neurons. Next, the image processing algorithm extracted neurodegeneration metrics used for sorting. One of the limitations of the image processing algorithm is that all four CEP dendrites must be visible, such that no dendrites are overlapped in the MIP. With standard imaging techniques where worms are immobilized on agar pads, nearly 50% of the worms are oriented such that their CEP neurons overlap in z-stack images. The microfluidic device design proved advantageous as a small overhead step at the end of the imaging channel (**Fig. S1**) resulted in an improved worm orientation that allowed that all four CEP dendrites were visible in the MIP (San-Miguel et al., 2016). After morphological analysis, worms were expelled from the imaging chamber into either a collection or trash outlet. We used this platform to sort worms into three groups based on dendrite morphology: least damaged (labeled “healthy” throughout this work), high dendrite breakage, and high bleb counts. Since most worms exhibited neurodegeneration, worms were classified as healthy if their four CEP dendrites had a median dendrite loss of less than 15% and less than 16 blebs. Worms were sorted into the blebs group if their dendrites had a median dendrite loss of less than 15% and more than 16 blebs. Worms sorted into the breaks group had a median dendrite loss greater than 15%.

The neurodegeneration metrics for each sorted group can be seen in **Figs 2B-D**. As expected based on the sorting criteria, healthy worms had the fewest breaks and blebs. Moreover, the breaks group contained significantly more breaks compared to the blebs and healthy groups, while the blebs group had significantly more blebs than the healthy and breaks groups. Interestingly, when comparing bleb count per dendrite length, there is no significant difference between the breaks and blebs groups. Animals with greater dendrite loss have fewer total blebs, but a similar bleb density as animals with mainly breaks, which suggests that these groups may have similar extents of neurodegeneration.

### d. Hallmark degeneration markers are associated with changes in DA-SWIP rates

After sorting worms into three distinct groups and allowing an overnight recovery, we measured their paralysis rate using the DA-SWIP assay discussed previously (**Fig. 2E**). Although there are no significant differences in the paralysis curves between the healthy, breaks, and blebs groups at early time points, differences between groups become apparent at later time points. After 8 minutes in 300 µM DA, nearly 50% of the healthy group are moving, while only 38% and 25% of the breaks and blebs group are moving, respectively (**Fig. 2F**). This difference is enhanced at 10 minutes, when 55% of healthy worms continue moving while only 20% of the breaks group and 25% of the blebs group is moving (**Fig. 2G**). Throughout all time points the breaks and blebs groups experienced similar paralysis rates, suggesting that blebs and breaks are representative similar levels of dopaminergic dysfunction.

### e. Summary

Here, we present a modified version of the swimming-induced paralysis assay by adding small amounts of DA. We demonstrate that DA-SWIP can reveal behavioral differences between known DA-deficient and DA-abundant strains from wildtype worms. Since worms exposed to 6-OHDA exhibit high rates of morphological dopaminergic degeneration, we hypothesized that these worms would produce less DA and behave more similarly to the DA-deficient *cat-2* worms. Interestingly, this was not the case. Worms exposed to 6-OHDA paralyzed at a quicker rate than wildtype worms, potentially indicating that 6-OHDA exposure results in altered dopamine signaling by affecting DAT-1 dopamine reuptake transporter or DOP-3, a D2-like receptor (Hardaway et al., 2015; Nass et al., 2002). We further asked whether hallmark neurodegeneration markers, such as breaks or blebs, resulted in differences in dopaminergic function in animals exposed to the same environmental (chemical) conditions.. We sorted over 1,000 worms into one of three collection groups or a trash outlet. We collected approximately 100 worms with healthy dendrites, dendrites with breaks, and dendrites with blebs. Subsequently, we measured the dopaminergic function of each group using DA-SWIP. Worms with healthy CEP dendrites paralyzed less than worms with either blebs or breaks, providing evidence that hallmark morphological neurodegeneration markers are indeed associated with neuronal dysfunction. However, it is intriguing that no difference was observed between blebs and breaks, suggesting the loss of function detected occurs with what has previously been regarded as a less severe form of damage (Bijwadia et al., 2021). The relationship between when dendritic damage first correlates with functional deficit should be further investigated to pinpoint the molecular changes that underscore loss of neuronal function.

## 4. Acknowledgements

Some strains were provided by the *Caenorhabditis* Genetics Center, which is funded by NIH Office of Research Infrastructure Programs (P40 OD010440).

## 5. Competing interests

No competing interests declared.

## 6. Author contributions

Conceptualization: ASC, KM, JM, ASM; Methodology: ASC, JH; Validation: ASC, JH; Formal Analysis: ASC, ASM; Investigation: ASC, KM, JH, JM, ASM; Writing: ASC, JH, ASM, KM, JM; Visualization: ASC, JH, ASM; Supervision: JM, ASM; Project Administration: JM, ASM; Funding Acquisition: JM, ASM.

## 7. Funding

National Science Foundation, Division of Molecular and Cellular Biosciences, 1947498 – Adriana San Miguel U.S. Department of Defense, W81XWH-18-1-0701 – Adriana San Miguel National Science Foundation, Division of Integrative Organismal Systems, 1838314 – Dr. Adriana San Miguel National Institute of Environmental Health Sciences, P42ES010356 – Joel Meyer National Institute of Environmental Health Sciences, R01ES034270 – Joel Meyer National Institute of Environmental Health Sciences, T32ES021432 – Katherine Morton

## 8. Data availability

Data will be made publicly available through a Dryad repository.

## Notes

### Competing Interest Statement

The authors have declared no competing interest.

